# Defining the transcriptional landscape of infiltrating immune cells in human and mouse bladder cancer

**DOI:** 10.1101/2020.07.09.195685

**Authors:** Haocheng Yu, John P. Sfakianos, Li Wang, Jorge Daza, Matthew D. Galsky, Nina Bhardwaj, Olivier Elemento, Bishoy M. Faltas, Jun Zhu, David J. Mulholland

## Abstract

The tumor immune cell landscape is composed of infiltrating myeloid cells (TIMs) and lymphocytes (TILs) that are important for regulating tumor progression and response to immune therapies. Since at least 70% of bladder cancer patients are poorly responsive to immune checkpoint blockade, there is a need for in depth analysis of the immune landscape both within individual tumor samples and between patients. To facilitate this, we have conducted single cell RNA sequencing (scRNA-seq) to map and transcriptionally define immune cells from 10 human bladder tumors. While all human tumors showed the of presence of major immune cell types, significant differences were observed in the relative numbers of these populations. To determine the translational utility of these findings we also performed scRNA-seq of TIMs-TILs isolated from mouse bladder tumors that were induced by the BBN carcinogen. In human and mouse comparison, we identified significant cross-species conservation in gene expression programs for NK, T, dendritic cells, monocytes and macrophages immune cell populations. Our data identified the conserved expression of key immunotherapy targets between human and mouse tumors including coordinately high gene expression of *PD-1*, *CTLA* and *IDO1* while also noting species divergent expression of *TIM3*, *LAG3* and *CD274* (PDL-1). To identify predictive immune cell-tumor interactions we computed conserved ligand-receptor (cell-cell) interaction scores between immune cell subsets and the epithelial tumor compartment. Together, this study defines the transcriptional status of bladder cancer immune cells and provides the rationale for future studies related to treatment response and therapeutic resistance.

## INTRODUCTION

Therapies targeting T cell function have changed the treatment management of non-muscle invasive, invasive and metastatic bladder cancer. This is well exemplified with PD-1/PD-L1 blockade which has shown clinically durable responses for patients who are cisplatin ineligible or with cisplatin-resistant metastatic bladder cancer [1–4][1–4]. However, only about 30% of patients experience a response that is clinically durable [5]. For these reasons, characterizing the state of treatment naïve tumor infiltrating myeloid cells (TIMs) and tumor infiltrating lymphocytes (TILs) is critical for the understanding of how immune cell transcriptional profiles may vary (1) between immune populations within individual tumors, (2) between tumors from different patients and (3) whether immune cell transcriptional programs are conserved between species. Such analyses may serve as a prerequisite to understanding signature differences that occur in tumors with *de novo* and acquired resistance as well as in the design of rationale immune based drug combinations.

Murine models of cancer serve an important role for the preclinical study of human tumor progression and identifying targets and rational drug combinations that are potentially effective in the clinical setting. Murine models that incorporate mutational landscape and lineage subtyping known to be present in human bladder tumors are limited. The N-butyl-N-4-hydroxybutyl Nitrosamine (BBN) carcinogen shares similar carcinogenic properties to tobacco products, which are implicated in more than half of bladder cancer diagnosis [6][5] [6]. In mice, BBN can induce the formation of invasive bladder cancer [7][7] and, like human disease, is composed of a heterogeneous array of mutational alternations [6] [7]. In-depth analysis of the molecular alterations in BBN tumors including whole exome and RNA sequencing have illustrated important similarities and differences between mouse and human bladder tumors [8] [7] [8]. Such mutational analyses have showed similarities in driver mutations, including *Trp53, Kmt2c* and *Kmt2d*. Despite this, there is a relatively poor understanding as to the state of infiltrating immune cells that occupy BBN tumors. Previous studies using human and mouse lung cancers [9] [8][9], have identified conserved and non-conserved transcriptional signatures in myeloid and accessory cells. Such analysis in bladder will provide a much-needed understanding of bladder TIMs-TILs and a platform to study specific immune lineages and the potential for translation to the clinical setting.

In this study, we have conducted scRNA sequencing analysis of immune cells isolated from human bladder tumors of varying pathological states. We highlight signature similarities and differences in myeloid and lymphocyte cell populations and implications for targeted therapies. We have conducted cross species analysis with the frequently used BBN murine bladder cancer model to identify conserved expression patterns and to qualify the applicability of this model towards preclinical testing of therapeutics that may modulate the CD45-postive tumor component. Finally, we argue that while important, expression signatures must be interpreted in the context of functional cell-cell interactions and intra tumoral immune cell localization.

## RESULTS

### Single Cell RNA-Sequencing of tumor cell populations in human and mouse bladder cancers

We aimed to define the immune cell landscape in human and mouse bladder tumors. To do this we obtained fresh bladder tumors from 10 patients including TURBTs (n=9) and cystectomy (n=1) **(Table 1)**. Patients (PT) with resected tumors were between 55 and 87 years old (median 73), contained a range of clinical stages, pT0-pT4 and were all treatment naïve. The one pT0 tumor was from a restaging TURBT on a non-muscle invasive bladder cancer patient. To generate mouse bladder tumors, FVB/NJ mice were exposed to the BBN carcinogen for 14 weeks, followed by 4 weeks of progression and subsequent tumor resection (n=3 tumors) [6] [7] [10] [11]. To prepare tissues for scRNA-seq, human and mouse tissues were enzymatically and mechanically digested to the single cell level, sorted using flow cytometry for CD45-positive and CD45-negative cells followed by rapid processing using the 10x Chromium platform. Sequencing was done on a Nextseq 500 platform and analyzed using Cell Ranger V2. Single cell data was filtered to exclude putative doublet populations, dead or mitochondrial high cells. Estimate numbers of total cells sequenced ranged from 807-1,927 for 10 human samples and 1,354-8,839 for 3 mouse tumors (M8524, M8525, M4950). The quality of cells was assessed by the number of genes detected, or percentage of sequence reads mapped to mitochondrial genes **(Fig. S1)**. Specific cell numbers before and after QC both for CD45 negative and CD45 positive cell sorts were also determined **(Table 2)**. For mouse, CD45-positive cells analyzed included 1,338 cells (4950P), 2,624 cells (8524P cells) and 4,938 cells (8525P). Mouse CD45-negative samples included 3,104 cells (4950N), 5,825 cells (8524N), 5,143 cells (8525N) and 5,618 cells (3928T) for analyses (where N = CD45 negative primary tumor cells, P = CD45 positive primary tumor cells, T = CD45 negative subcutaneous tumor cells).

### Comparative analysis of CD45-negative populations in mouse and human bladder tumors

We performed clustering analysis to identify distinct cell types in CD45 negative fractions in human and mouse bladder tumors. In both, we identified four cell types including epithelial, endothelial, fibroblasts and myofibroblasts **(S2A-B)**. Cell types are annotated by highly expressed marker genes including *KRT* (epithelial), *PLVAP* (endothelial), *MMP2* (fibroblasts) and *RGS5* (myofibroblasts). Both human and mouse epithelial cells showed high expression of *PSCA-Psca*, *KRT7-Krt7* and *KRT19-Krt19*. Epithelial cell-specific genes that were highly expressed in mouse tumors included *Ly6d*, *Krt14*, *Gsto1*, *Sprr2a3*, *Wfdc2*, *Krt15*, *Sfn*, *Fxyd3* and *S100a14* while in human cells *SPINK*, *HPGD*, *CLDN4*, *RARRES1* and *CD24* were highly expressed. In mouse and human tumors, fibroblasts were concomitantly high for *COL1A-Col1a*, *COL1A2*-*Col1a2*, *COL3A1*-*Col3a1* and *DCN*-*Dcn*. Myofibroblasts were both high for RGS5-Rgs5 and *HIGD1B*-*Higdlb.* We further clustered epithelial cells to examining variability in transcriptional profiles among sample types and tumor source. Examining sample type and tumor source, we determined that epithelial cells in mouse primary and subcutaneous (SQ) tumors formed distinct clusters **(Fig. S2C)**.

Epithelial cells from individual human tumors formed distinct clusters **(Fig. S2C, right)** suggesting the presence of transcriptional diversity amongst human tumors. Interestingly, when clustering mouse and human CD45 negative cells, epithelial cells in human tumor 171 were more similar to ones in mouse 3928N while epithelial cells in human 2a were closer to ones in mouse 8524N and 8525N **(Fig. S2D) (Supplementary Data 1)**. Subcutaneous tumors were characterized by high expression of *Spp1* (osteopontin), *Lgals7*, (intracellular galectin-7), *Ggct* (GGCT), *Pthlh* (PTHLH), *Slpi* (SLPI), *Igfbp4* (IFBP4) as well as reduced expression of markers of differentiation (*Krt5*, *Krt13*, *Krt15*) **(Fig. S2B)**. Interestingly, by comparing epithelial cell gene expression in the primary and SQ tumors, we observed that SQ tumors (clusters 0+4) showed higher activity of Myc signaling. Genes expressed higher in primary mouse tumors were enriched in pathways related to TNF alpha and hypoxia, apoptosis, xenobiotic metabolism, adipogenesis, oxidative phosphorylation and EMT **(Supplementary Data 2)**. *Together, these data indicate that human bladder tumors have diverse CD45 negative transcriptional profiles while mouse primary BBN tumors are more similar to each other but divergent from SQ tumor profiles. Furthermore, human and mouse primary BBN tumors share intrinsic CD45 negative transcriptional signatures.*

### Clustering analysis of mouse and human bladder tumor infiltrating immune cells

CD45-positive cells in mouse and human tumors were partitioned into cell clusters **(Fig. 1A)**. To annotate human cells we used whole transcriptome profiles of FACS sorted subpopulations while for mice we used data sets from the IMMGEN consortium [12] [9] [10] [11]. Based on highly expressed genes, immune cell subsets were annotated as: T and NK (T-NK) cells (*CD3E*, *Cd3e*), macrophages + monocytes (*LYZ*, *Lyz2*), classic DCs (*CCR7*, *Ccr7*), pDCs (*TCF4*, *Tcf4*), mast cells (*MS4A2*, *Ms4a2*), B cells (*MS4A1*, *Ms4a1*), plasma cells (*MZB1*, *Mzb1*) and neutrophils (*CD24A*, *Cd24*) **(Fig. 1B, C)**. We identified all major immune cell lineages including myeloid cells (mast cells, neutrophils, classic DCs and plasmacytoid DCs) and lymphoid cells (T cells, NK cells, B cells and plasma cells). Curiously, we did not detect a cluster of mouse plasma cells distinct from B cells.

**Fig. 1.**
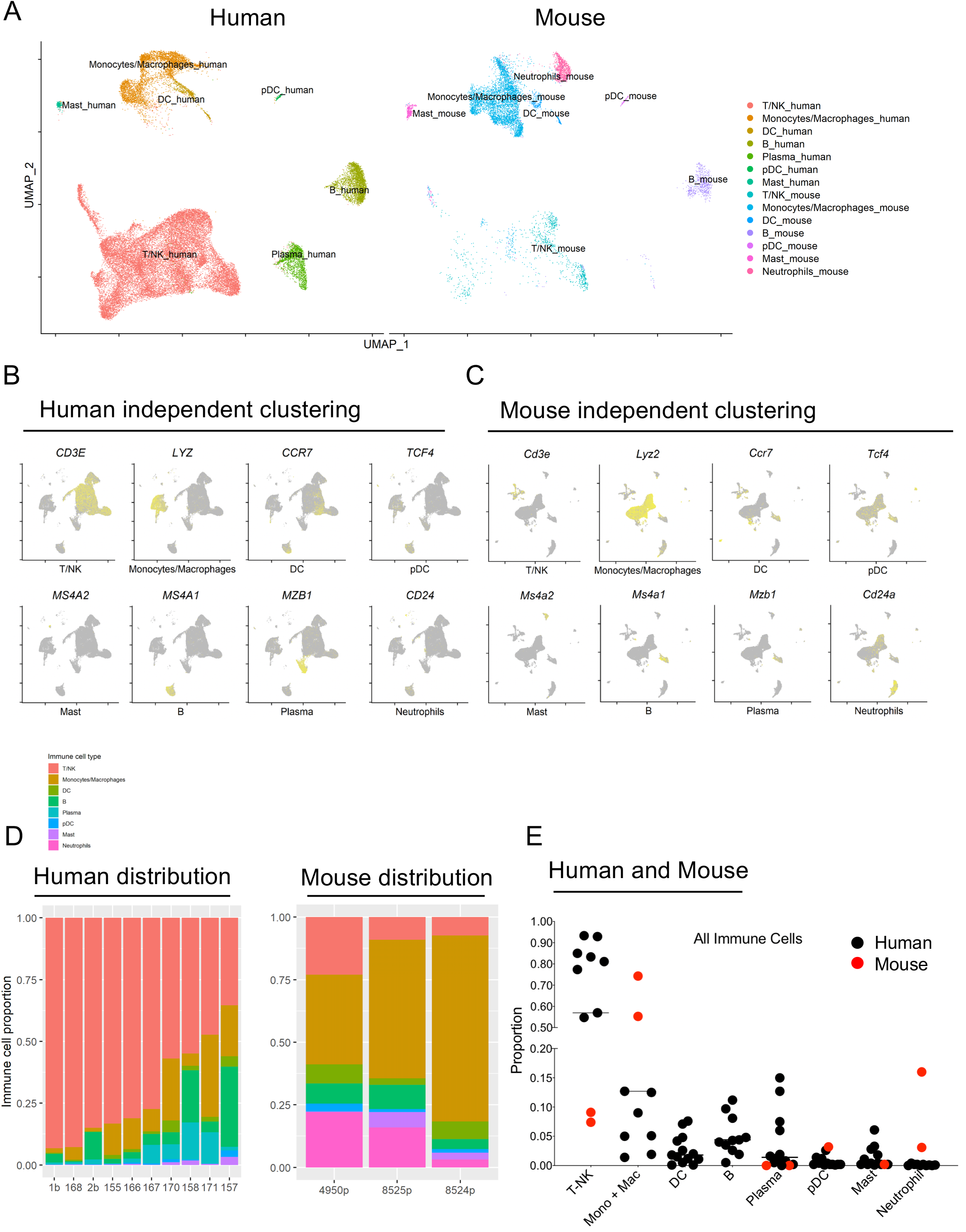
Single cell transcriptional profiling of human and mouse bladder tumor infiltrating immune cells. (A) UMAP plots clustering for immune cell types in pooled human (n=10) and mouse (n=3) tumors. UMAP plots showing genes that define immune cell types in (B) human and (C) mouse TIMs. Immune cell proportions for individual (D) human and mouse bladder tumors. (E) Proportion of immune cell populations in CD45-positive cells in human and mouse bladder tumors.

Next, we searched for differences and similarities in the abundance of major immune populations in human and mouse tumors. In all human bladder tumors, the predominant immune cell types were T cell and NK cell (T-NK cells) while in all mouse tumors monocytes and macrophages (monocyte-macrophage) cell derived populations predominated **(Fig. 1D)**. Despite these trends, we observed considerable patient-to-patient variation in the composition of immune cell populations. For example, we detected the presence of T-NK “higher” (1b, 168, 2b, 155, 166, 167, >75% CD45+ of cells in composition) and T-NK “lower” (158, 170, 171, 157, <55% CD45+) human bladder tumors. In mice, all tumors could be considered T-NK “low” while monocyte-macrophage fractions were “high” as compared to human tumors. Of 10 PTs, we identified 3 tumors (170, 171, 157) with comparable macrophage-monocytes fractions to mouse tumors **(Fig. 1D&E)**. Of 10 tumors assessed, 157 was staged as a T0 and showed the least proportion of T-NK cell infiltration and the greatest content of B cells. Despite this trend, the degree of T-NK infiltration did not show a clear association with the clinical cancer stage or source of tissue (cystectomy or TURBT) **(Fig. 1D) (Fig. S3) (Table 1)**.

### Conservation in NK and T cell transcriptional programs between mouse and human bladder tumors

To conduct more detailed analysis, we partitioned T-NK cells into 8 clusters in human and 6 clusters in mouse **(Fig. 2A&B)**. Clusters were annotated as CD4 T cells defined by *CD4-Cd4* expression, regulatory CD4 T cells by *FOXP3-Foxp3*, naïve-memory CD4 T cells by *CCR7-Ccr7*, CD8 T cells by *CD8A-CD8a*, NK and effector CD8T cells by *NKG7-Nkg7*, or exhausted CD8 T cells by *PDCD1-Pdcd1* **(Fig. 2C&D)**. In total, there was 1 cluster of NK cells (mouse, human), 5 clusters of CD4 T cells (mouse, human), 1 cluster of CD8+ T cells (mouse) and 3 clusters of CD8+ T cells (human) **(Fig. 2A-D)**. We calculated correlations among average expression profiles of individual cell clusters using 2000 highly variable genes among all cells and assessed mouse and human T-NK cells for functional similarities in all T-NK cell clusters **(Fig. 2E, Supplementary Data 3)**. The hierarchical analysis showed co-clustering in (1) mouse-human CD8+ T cells (exhausted, effector), (2) NK cells, (3) mouse-human CD4+ T cells (exhausted, regulatory) and (4) human CD4+ T cell (naïve, memory) **(Fig. 2E)**. Using heat map alignments between species, we observed multiple clusters of commonly overexpressed genes between mouse and human that were most notable in exhausted CD4 T cells (set 1), regulatory CD4 T cells (set 2), NK cells (set 3) and exhausted CD8 T cells (set 4) **(Fig. S4A)**suggesting the possibility of functional similarities between species including particular genes constituting these immune cell populations as shown in **Fig. S4B**. The conservation in mouse and human T-NK cell programs prompted us to identify functional similarities related to cell differentiation (CD), naïve status, inhibitory function, effector function, co-stimulatory function, transcription factor expression and T regulatory function **(Fig. 2F). We detected** coordinately high expression of many signature genes between mouse and human for exhausted CD8+ T cells populations. Examples of such genes related to cell differentiation (*CD8A-Cd8a*, *CD8B-Cd8b*), inhibitory receptor response (*LAG3-Lag3*), cytokine-effector molecules (*GZMA-Gzma*, *GZMB-Gzmb*, *NKG7-Nkg7*), transcription factors (*EOMES-Eomes, ID2-Id2*) and Treg markers (*FOXP3-Foxp3*). Other immune cell populations including NK cells, CD4+ regulatory T cells and CD4+ exhausted T cells also showed concordant high expression of genes in multiple functional classes **(Fig. 2F)**. Despite these qualitative similarities, we also identified several obvious differences between T-NK functional populations between mouse and human. While human tumors showed a higher fraction of T-NK cells **(Fig. 1D)**, mouse tumors were higher in DCs, pDCs, B cells, Mast and Neutrophils **(Fig. S3C)**. BBN mouse tumors also showed higher proportions of naïve, regulatory and activated CD4 T cells and “double negative” (DN) T Gamma Delta among T-NK populations **(Fig. 2F) (Fig. S3A)**. Interestingly, double negative immune cells were not only low for CD4 and CD8 markers were also low in expression of effector molecules including *Il2*, *Gnma*, *Prf1*, *Gnmb*, *Gzmk*, *Ifng* and *Nkg7* (green panel, Cytokines and effector molecules) **(Fig. 2F)**.

**Fig. 2.**
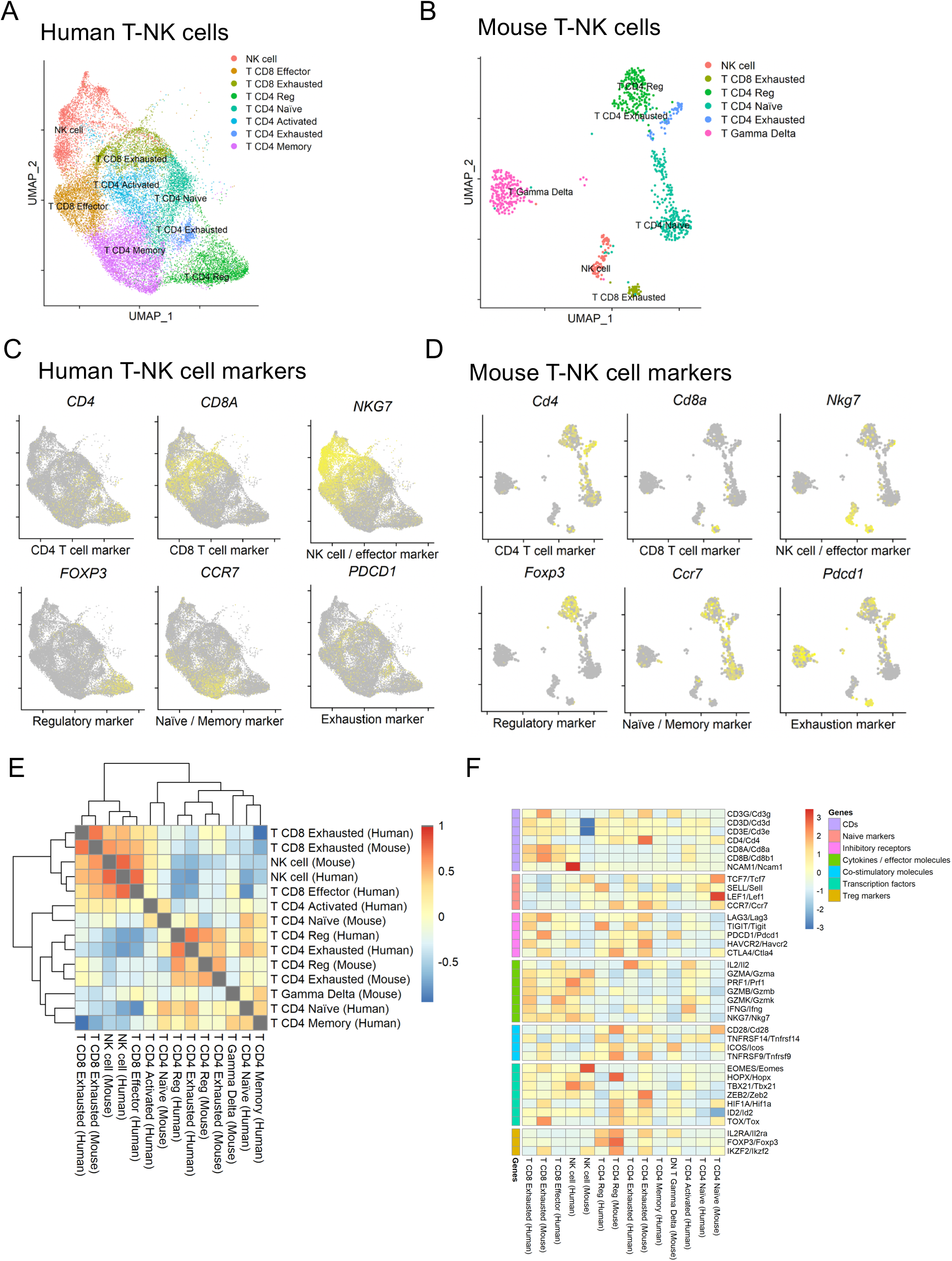
Identification of NK cell and T cell gene expression programs in human and mouse bladder TILs. Integrated UMAP analysis for (A) human and (B) mouse T-NK cells. Expression of T-NK specific markers in pooled (C) human and (D) mouse bladder tumors. (E) Correlation expression between mouse and human T-NK cells. (F) Comparison of functional marker expression for T-NK cells between human and mouse. See **Supplemental Data 3**.

Collectively, human bladder tumors and the BBN induced mouse bladder tumors share T-NK cell transcriptional programs including functional markers related to T cell activation, exhaustion and regulatory function.

### Species and organ conservation in macrophage-monocytes populations in mouse and human bladder cancers

When clustering macrophage-monocytes, we identified 13 cell clusters in human bladder tumors and 15 cell clusters in mouse bladder tumors, respectively **(Fig. S5)**. We annotated these clusters using established gene signatures for monocyte and macrophage populations [13, 14][12–14], and merged the 13 clusters into 8 functional clusters for human samples including 3 monocytes clusters with high expression of monocyte signature genes (*VCAN*, *FCGR3A*, *CXCL8*), 4 macrophage clusters (*CD163*, *C1QC*, *CXCL11*, *SPINK1*) and one myeloid-T cell positive cluster (*CD14*, *LYZ*) **(Fig. 3A)**. For mouse tumors, we merged 13 clusters into 10 functional clusters including 3 as monocytes (*S100a9*, *TremI4*, *Vcan*), 5 as macrophages (*Cd74*, *Cdh1*, *Ctsd*, *Gpnmb*, *Lyve1*), one myeloid DC (*Ccr7*, *Cd74*, *MhcII*) and one myeloid-T cell positive **(Fig. 3B)**. The *S1009a* population was identified to be similar with both species having high expression of *S100a9*, *S100a8* and *Vcan* **(Fig. 3C, D)**. These analyses were conducted the 2000 most highly variable genes among cells. Functional similarity was assessed among clusters by calculating correlations between average profiles of individual clusters **(Fig. 3E, Supplementary Data 4, Supplementary Data 5**). The data indicate that human monocyte and macrophages have more organized clustering while in mice clusters appeared more complex and in greater numbers including those not identified in human tumors **(Fig. 3E)**. When comparing clusters by alignments we identified at least 6 sets of commonly over expressed genes including human and mouse myeloid T cells (sets 1 and 2), mouse *S100a9* monocytes and human *CXCL8* monocytes (set 3), mouse *Ctsd* macrophages and human *SPINK* macrophages (set 4), mouse *Cd74* macrophages and human *CXCL11* macrophages (set 5) as well as mouse *Gpnmb* macrophages with human *C1QC*-*CD163* macrophages (set 6) **(Fig. S6A)**. *These data identify the presence of highly co-expressed genes between mouse and human macrophage-monocyte populations.*

**Fig. 3.**
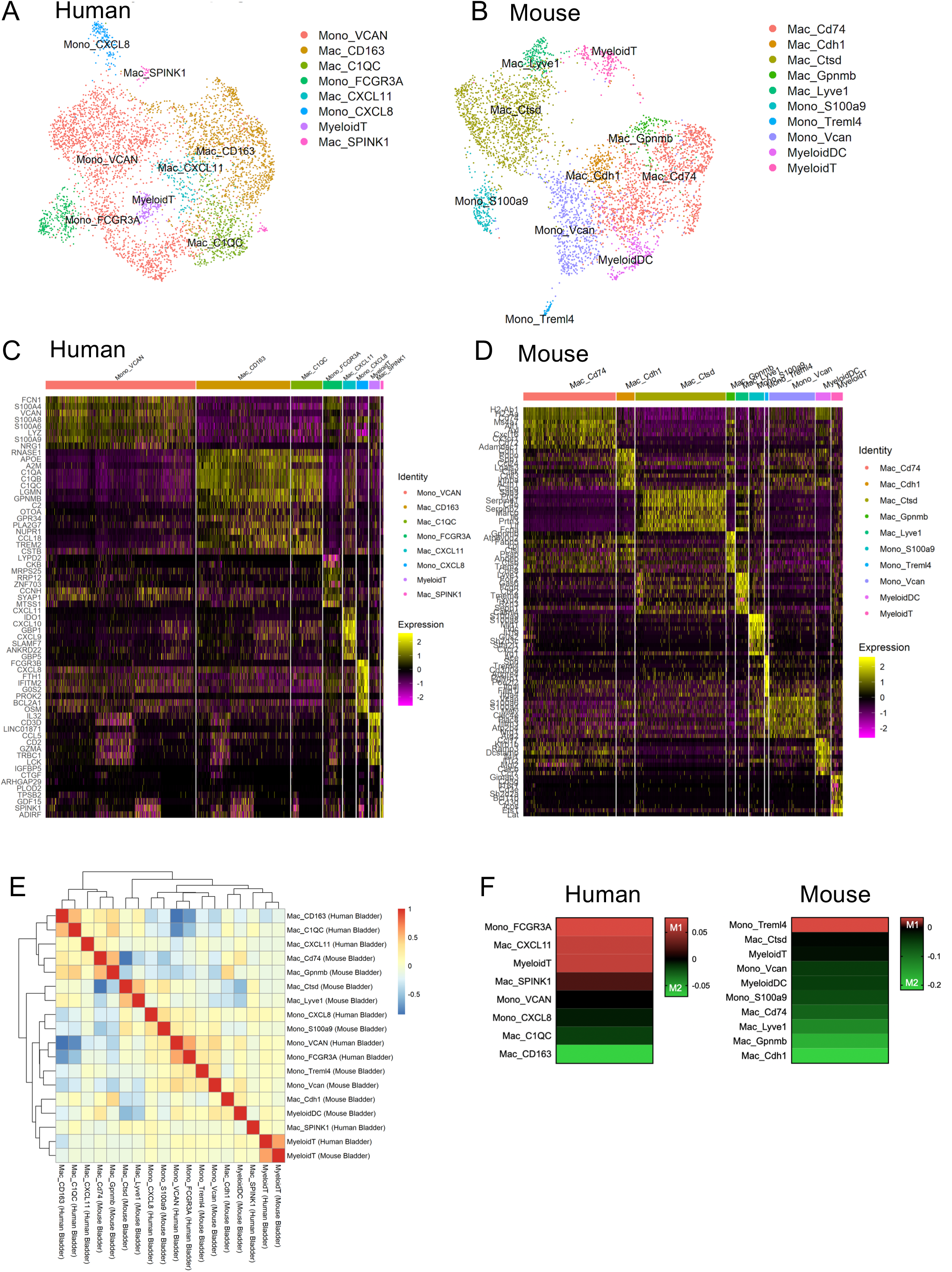
Bladder tumor monocytes and macrophages show species and organ conservation. Integrated UMAP analysis for (A) human and (B) mouse monocytes and macrophages. Heat maps showing major gene expression profiles for monocyte-macrophages derived from (C) human and (D) mouse bladder tumors. (E) Correlation heatmap between human and mouse immune cell populations. (F) Heat maps showing expression of M1 and M2 genes for human and mouse monocytes-macrophages. See **Supplemental Data 4-9**.

### Conservation in gene expression between tumors from different organs

With the observed conservation between human and mouse monocytes-macrophages, we evaluated whether such similarities in functional programs could exist with other organs. Thus, we clustered monocytes-macrophages from human lung and human bladder using the established signature genes for lung tumors [9] **(Fig. S6B)**. We identified at least 7 sets of significant transcriptional overlaps in human bladder and lung tumors including bladder clusters *C1QC* (sets 1, 2), *CXCL11* (set 3), *CD163* (set 4) as well as bladder monocyte clusters with high *CXCL8* (sets 5, 6) and *VCAN* (set 7). When we compared mouse lung *versus* mouse bladder, we observed even greater conservation including at least 13 sets of common highly expressed signature genes between bladder monocytes with lung monocytes (sets 1-3) and bladder macrophages with lung macrophages (sets 4-13) **(Fig. S6B, lower)**. Interestingly, we found certain monocyte populations to be present in bladder and lung of both species including classic CD14 positive monocytes as defined in human bladder *VCAN-Vcan* cells, mouse *Vcan*-*S100a9* cells, human lung clusters 1+3 and mouse lung clusters 1+3. CD16 positive monocytes were also identified in human bladder *FCGR3A* cells, human lung cluster 2, mouse bladder *TremI4* cells and mouse lung cluster 2. Evaluation of the identified populations was further assessed by clustering analysis showing major separations between monocytes and macrophages in human and mouse TIMs **(Fig. S6C)**. *Thus, these analyses show the presence of specific monocytes populations shared between species and different organs.*

### Pathway Enrichments

To further understand the relationship of monocyte and macrophages populations, we conducted pathway analysis on individual cell populations in human and mouse samples **(Supplementary Data 6, 7)**. To do this we compared module scores for each pair of human or mouse immune cell subclusters to get rank-sum test p-values, thus determining whether there is a difference of pathway activation between any cluster pair. In humans, we observed monocytes (*VCAN*, *CXCL8*) with activation of multiple pathways including MYC and E2F target **(Fig. S7A) (Supplementary Data 8)**. From here we conducted comparisons of major signaling pathways including MYC, PI3K-AKT, interferon-gamma and TNF-alpha between each pair of different monocyte-macrophage populations **(Fig. S7B)**. In these analyses we identified several outlier populations including human monocyte (*VCAN*) and myeloid T cell populations that were high in MYC V1-V2 (heat map 1) and PI3K-AKT signaling (heat map 4), as compared to any other populations. We identified Mac (*CXCL11*) cells to be high in interferon gamma response (heatmap 7) but low in activation of other pathways. Conversely, in mouse, Myeloid T cells showed low activation of all four signaling pathways (heat maps 2, 5, 8, 11) while Monocyte (*Vcan*) cells also demonstrated high activation of PI3K-AKT signaling (heat map 5) **(Supplementary 9)**. *Together, these data show that monocyte-macrophages populations show differential signaling dependencies.*

### Defining M1 + M2 macrophages signatures

Macrophages and monocytes are generally defined as having an M1 and or M2 transcriptional profile with pro-inflammatory and anti-inflammatory functions, respectively. To understand the M1-M2 status here we assessed human and mouse cell clusters based on expression of genes that could define monocyte-macrophage populations as either M1 (e.g. *CXCL9*, *IDO1*) or M2 (e.g. *CD163*, *CCL18*) [15][12] [13] [14]. We compared M1 and M2 module scores for each clusters based on expression of M1 or M2 module genes by the rank-sum test **(Fig. 3F) (Supplementary Data 11, 12)**. Human pathway p value calculations indicated that human *FCGR3A* monocytes, *CXCL11* macrophages and Myeloid T cells were more M1-like (with higher M1 module scores than M2 module scores) while *VCAN*, *CD163*, *C1QC* and *CXCL8* defined cell clusters were more M2-like. In mouse, with the exception of TremI4 defined monocytes, all populations were M2-like **(Figure 3F)**. *These results also show that, compared to human, mouse monocyte-macrophage populations cluster into more transcriptionally defined populations. The data supports that monocyte and macrophages signature conservation occurs not only between mouse and human bladder cancer cells but between different organs of the same species. Pathway analyses indicates that M2-like cells predominate particularly in mouse bladder tumors.*

### Conservation in expression of major therapeutic targets between human and mouse T-NK and monocytes-macrophages populations

Investigating the immune cell landscape in mouse and human bladder tumors provided the opportunity to evaluate the transcriptional status of therapeutic targets. We considered the expression status of a panel of genetic targets catalogued from genetically engineered mouse models as well as genes that encode proteins targeted by immunotherapeutics in CD45-positive bladder tumor cells [8][9] **(Fig. 4)**. In doing so, we identified examples of conserved genetic targets between mice and humans relevant to different immune cell types **(Fig. 4A)**. These included coordinately high expression of genes in human and mouse T-NK cells (*FOXP3*, *LCK*, *CD3E*, *INFG* and *CD2)*, monocytes-macrophages (*CTSK*, *MAFB*, *ITGAM*, *CD68*, *MERTK*, *CSF1R*, *CEBPA* and *NR4A1*), DCs (*BATF3*, *XCR1*, *ZBTB46*, *CLEC9A* and CD207), pDCS (*FLT3*, *CXCR3*) and mast cells (*CPA3*, *CMA1)*. While many gene target-cell type associations conserved well between mouse and human cells, we identified clear differences in multiple immune cell populations including: high *CD8A* expression in human T-NK cells but low in mouse, high *CCR2* expression in human monocytes-macrophages but low in mouse, high *S100A8* expression in human monocytes-macrophages but low in mouse. Genes that were <u>high in mouse</u> but low in human tumor-associated immune populations included high *NR4A1* expression in mouse neutrophils, high *Cd4a* in mouse T-NK cells, high *Cd8a* in pDCs and high *Cx3cr1* in mouse mast cells **(Fig. 4) (Fig. S8)**.

**Fig. 4.**
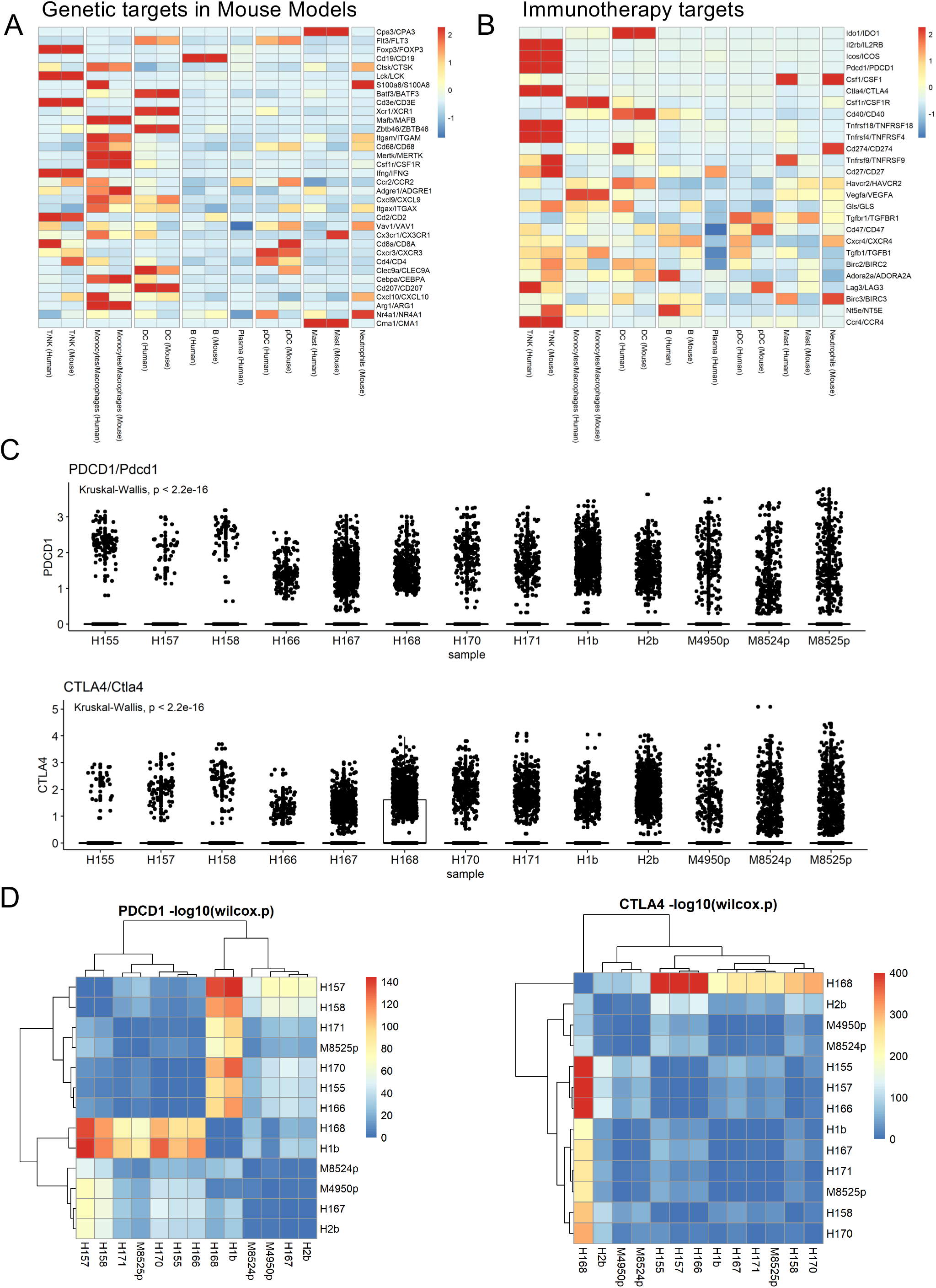
Species conservation of genetic and immunotherapy targets in bladder tumors. Clustering for similarities between human and mouse bladder immune cells using (A) genetically defined targets and (B) clinical immunotherapy targets. (C) Single cell gene expression of immune checkpoint markers (PD1, CTLA) in independent mouse and human bladder tumors (Kruskal Wallis generated p values). Horizontal lines represent 1^st^, 2^nd^ (media) and 3^rd^ quantiles of the data. (D) Clustering analysis showing similarity in *Pdcd1-PDCD1* and *Ctla4-CTLA4* expression between human and mouse bladder tumors. Those having greater red expression are more divergent according to Wilcox p values.

To understand the relative expression of immunotherapy targets in human and mouse immune cells, we assessed a panel of known targets including those with clinical implications [9]. In human and mouse NK-T cells we observed high coinciding expression of the following immune checkpoint proteins, *Icos-ICOS*, *Ctla4-CTLA4*, *Pdcd1-PDCD1* as well as others associated with T cell activation and response including *Il2rβ-IL2RB*, *Tnfrsf18-TNFRSF18*, *Tnfrsf4-TNFRSF4*, *Cd27-CD27* and *Ccr4-CCR4*. In some instances high levels of immune checkpoint proteins were observed in non T-NK immune cells including *Havcr2-HAVCR2* in human and mouse DC cells **(Fig. 4B)**. Identifiable species differences included higher *LAG3* levels in human mouse T-NK cells. Interestingly, the immune suppression molecule *Nt5e-NT5E* (CD73) [16, 17] was expressed higher in mouse than human NK-T cells and higher in human than mouse B cells. Other genes that were differentially expressed between species were identified in pDC cells, mast cells and neutrophils **(Fig. 4B)**.

With the variable levels of TIMs-TILs between individual human and mouse tumors, we considered the spectrum of gene expression for immune check point proteins in 10 patient tumors and 3 mouse tumors. Importantly, wide ranging expression of the *PDCD1*, *HAVCR2*, *CTLA4* and *CD274* (PDL-1) was observed amongst human tumors (**Fig. 4C, Fig. S9A)**– a finding compatible with the observed wide range in the relative numbers of immune cell types between individual tumors (low average expression = 166, high average expression = tumor 168, 1b, Kruskal-Wallis test p values <1e-15 for PDL-1 and CTLA4). We conducted Wilcox tests to identify similar and more divergent expression values between human-human, mouse-mouse and human-mouse tumors. With this analysis we observed the following trends: (1) tumors H168 and H1b were similar with respect to *PDCD1* and *LAG-3* expression (rank-sum test p-value<1e-50by compared H168 or H1b to any of other samples), (2) H168 was unique with high *CTLA4* expression (rank-sum test p-value<1e-50), (3) mouse tumors only clustered together exclusively for *CD274* (PDL-1) expression and (4) M8524 was unique amongst all tumors with respect to *HAVCR2* expression and (5) Tumors M8524 and M4950, clustered more closely to other human tumors than to the remaining mouse tumor (8525) **(Fig. S9B)**. *Together, these data show conserved expression of many genetically defined and major immune checkpoint proteins between human and mouse T-NK and monocyte-macrophages populations. However, these data also highlight the heterogeneity in target expression amongst individual human and mouse tumors.*

### Evaluating cell-cell interactions using established ligand-receptor interactions

Our analysis identified mouse-human species conservation in regulatory markers of immune cell activation and exhaustion. Since TIM and TIL function is regulated by cell-cell interactions including ligand-receptor conjugation, we leveraged a database of about 1800 known ligand-receptor interactions [18]. With an established computational approach [19], we clustered for heightened co expression of ligands and receptors in pooled mouse and human bladder tumors focusing on potential interactions between (1) tumor epithelia with T-NK cells and (2) tumor epithelia with macrophages-monocytes **(Fig. 5A)**. We calculated receptor expression and average ligand expression in the cell types of interest to determine calculated ligand-receptor interaction scores [19]. Cell labels shown in Fig 5A were written as (cell type expressing the ligand) – (cell type expressing the receptor) and were used to calculate the interaction scores represented in a heat map. Interactions having a score of greater than 4.0 across in paired cell-cell interactions are displayed as predicted interactions that were conserved between human and mouse tumor cell ligands and T-NK cells receptors including *PKM-CD44*, *B2M-CD3D* and *B2M-CD3G*. Putative epithelia ligand and macrophage/monocyte receptor interactions included *TIMP1-CD63*, *RPS19-C5AR1, FN1-CD44, FN1-C5AR* and *APP-CD74*. These findings have identified potential mechanisms of treatment resistance in the analyzed human and mouse tumors. For example, B2M or beta-2-microglobulin is an essential component of MHC class I antigen presentation and is enriched threefold in melanoma patients nonresponsive to CTLA4 or PD-1 treatment as compared to responsive patients [20]. *RPS19-C5AR1* interactions are important in promoting tumor growth and immune suppression by interactions with myeloid derived suppressor cells, leading to altered T cell response [21]. *TIMP1*, a secreted glycoprotein, is expressed at high levels in bladder cancer and has been demonstrated to interact with CD63 in glioblastoma [22]. When assessing interactions between immune cell ligands and epithelial receptors we detected species conserved T-NK ligand-epithelia interactions including *HMGB1-SDC1*, *CCL5-SDC1*, *CCL5-SDC4*. Macrophage-monocyte ligand-epithelial interactions included *TIMP1-CD63, THBS1-SDC4, THBS1-SDC1, THBS1-ITGB1, SPP1-ITGB1*, *HMGB1-SDC1, HBEGF-CD9, CD14-ITGB1*. SDC1 is an established risk factor for poor survival in bladder cancer [23]. *Collectively, these data show that the majority of predicted interactions involve cell-cell interactions between mouse tumor epithelia and macrophages-monocytes and that many of these interactions are conserved between human and mouse tumors.*

**Fig. 5.**
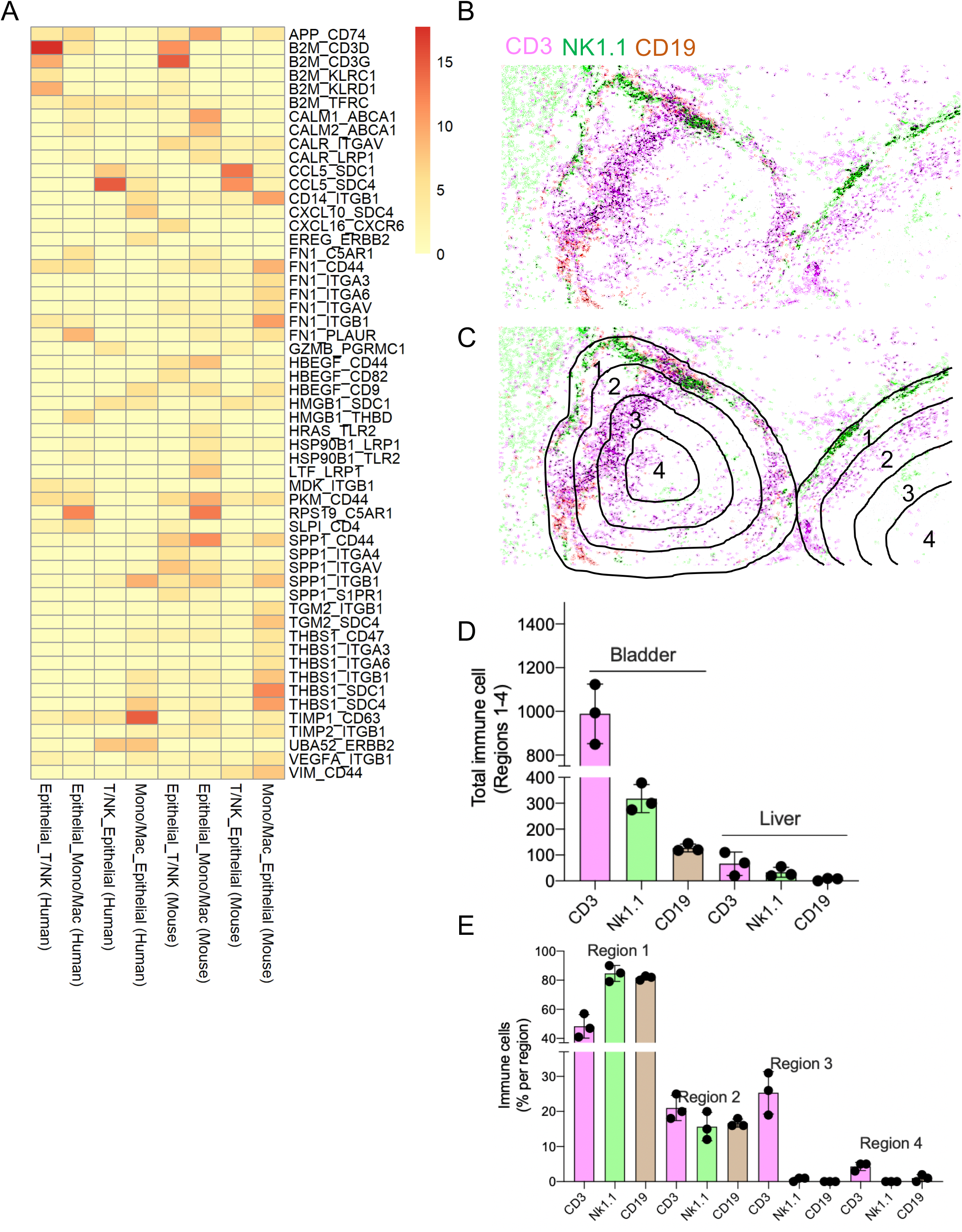
Determining cell-cell interactions and immune cell localization in human and mouse bladder tumors. (A) Heat map showing cell-cell interactions between immune cell populations (T-NK, macrophages and monoctyes) in pooled human (n=10) and mouse (n=3) bladder tumors. Cell type interaction scores are labelled as (cell type ligand expression) – (cell type receptor expression). Only interactions shown all have a score of >4. (B) Immunostains in mouse BBN induced bladder lesions for T cells (CD3+), NK cell (NK1.1+) and B cells (CD19+) shown as a pseudo color overlay. (C) Assignment of regions 1-4 based on equal distances from the lesion center to the lesion periphery. (D) Measurement of immune cell numbers in regions 1-4 in independent bladder and liver lesions. (E) Distribution of each immune cell type within each region of bladder lesions as a percentage of total immune cells found in regions 1-4.

### Myeloid and lymphocyte localization patterns complement transcriptional immune cell signatures

While single cell sequencing provides important information about the transcriptomic landscape of tumor immune cells, such analysis does not reveal information related to the cellular concentration (e.g. diffuse *versus* nested) and localization of immune cells (peripheral *versus* infiltrated). Immune cell localization studies may have significant implications for tumor progression and response to immune therapies. Thus, to extend our scRNA-seq analysis, we have applied a pathology-based approach to highlight the differences in absolute immune cell numbers and degree of infiltration to bladder lesions. Subsequent to propagating primary mouse bladder tumors and implants in the liver, tissues were resected, formalin fixed and processed for immunostaining using consecutive tissue section stains for CD3 (total T cells), NK1.1 (NK cells) and CD19 (B cells). Immunostains were visualized using DAB exposures, digitally scanned and converted to a monochrome signal. Unique pseudo colors were then applied to each DAB exposure in Adobe Photoshop imaging software followed by image superimposition **(Fig. 5B, Fig. S10)**. Independent images were then assessed using QuePath software for counting metrics that included (1) absolute immune cell numbers and (2) localization of each immune cell type with respect to the distance from the lesion center. Using concentric contour lines within the lesion we defined 4 regions spaced equally but with increasing distance from the lesion perimeter **(Fig. 5C)**. Compared to the liver microenvironment (not shown), primary bladder tumors contained significantly more T, NK and B cells. Of these, CD3 cells showed the highest absolute numbers. In terms of localization, NK and B cell were found predominantly near the lesion perimeter as defined by histological examination (region 1) while CD3+ T cells could be detected in significantly higher numbers in regions 2 and 3. In region 4, immune cells were present in 5% of less of total counts **(Fig. 5D, E)**. In bladder lesions propagated in the liver, nearly all immune cell were found in region 1 (tumor periphery). *Thus, using a pathology based multiplex counting approach we provide a cost effective method to provide qualitative and quantitative information about TILs-TIMs that may function in a complementary fashion to single cell transcriptomic analysis.*

## DISCUSSION

In this study we conducted a transcriptomic analysis of tumor infiltrating myeloid (TIMs) and tumor infiltrating lymphocyte (TILs) purified from human bladder tumors and carcinogen induced mouse bladder tumors. Our study aimed to (1) qualitatively and quantitatively analyze the immune landscapes contained in single cell tumor populations, (2) identify conserved and divergent transcriptomic signatures between human and mouse tumors, (3) determine whether the BBN carcinogen model shows maintenance of clinical targets of immune checkpoint inhibition, (4) apply a computational approach to understanding cell-cell interactions in tumor and (5) define a cost effective immune cell localization approached to help contextualize transcriptomic data.

Our analysis was leveraged by the isolation of purified (sorted) viable CD45 positive and negative cells in fresh human tumors associated with defined clinical, pathologic and demographic backgrounds. Single cell processing of purified cells and unsupervised clustering analysis lead to the profiling for human and mouse bladder immune cells. In both species, we use established cell type specific gene expression signatures to annotate individual myeloid and lymphocyte cell types. While the 10 human bladder samples assessed here showed marked variability in TIM and TIL proportions, the number of transcriptional clusters (or states) remained similar throughout.

The most obvious difference observed between human and mouse bladder tumors was the content of T-NK and monocyte-macrophage populations – where human tumors were composed of a markedly higher fractions of T-NK cells and mouse tumors with higher monocyte-macrophage fractions. High infiltration with TAMs (tumor associated macrophages) is associated with poor clinical outcomes in several types of cancer including breast, ovarian and lung [24, 25]. Thus, the high TAM levels in mouse tumors observed here may highlight the immunosuppressive state often observed in mouse tumors rapid progression kinetics as well as the homogeneous landscape of pathology as compared to human disease. Interestingly, pathway analysis showed that while human tumors were composed of a mixed population of M1-like and M2-like cells, mouse tumors were composed of predominantly M2-like monocytes and macrophages. These observations suggest that the BBN mouse model may be well suited for understanding how M2 populations promote immunosuppression and tumorigenesis. As well, TAMs are highly plastic and capable of conversion between different activation states including transitions between M1 and M2 types that may occur during organ development, cancer initiation and in response to therapy [10] [26]. The identified populations in our study may facilitate the study of these population transitions in future analysis.

Conservation in TIM and TIL signatures were detected not only between human and mouse bladder monocytes-macrophages but also between bladder and lung. These observations suggest that in both bladder and lung certain monocytes-macrophages populations may have general tumor specific (inflammatory) functions while resident macrophages may be more related to normal or organ specific functions. Future studies should attempt to discriminate transcriptionally signatures of macrophages in normal bladder from those assigned to bladder cancer. Additional areas of interest including understanding the effects of TAMs on therapeutic resistance including determining the effects of anti-cancer treatments on various subsets or transcriptionally defined clusters of monocytes-macrophages [27] [28].

Single cell transcriptomic analysis of CD45-negative cells revealed a marked separation of gene signatures for primary organs *versus* those maintained as early passage subcutaneous tumors. The propagation of subcutaneous tumors is typically done for practical reasons (ease of implantation, visualization of tumor growth). However, with the dramatic drift in expression signatures away from those of the primary organ, we argue that subcutaneous tumor signatures may not reflect expression signatures occurring in the majority of human tumors. Nevertheless, our study identified that some human tumors (e.g. tumor 171) preferentially clustered with mouse subcutaneous tumors. Thus, the choice of therapeutics testing in subcutaneous versus primary tumor mouse models should be done with careful consideration of the transcriptomic signature of the initial (donor) human tumor.

If the goal of defining immune cell states is to assign tumors as immune “cold” versus “hot” and to predict treatment response, then transcriptomic single cell analysis requires complementary assays that evaluate TIM and TIL intra tumor localization. Our study considered two approaches including a (1) computational cell-cell interaction scoring and (2) a cost-effective immunohistochemistry method for measuring immune cell infiltration on a quantitative and qualitative scale. Cell-cell interactions scores calculated from scRNA-seq data [19] identified several potential ligand-receptor interactions including those involving markers for tumor invasion (TIMP1) and resistance to immune checkpoint blockade (B2M or beta-2-microglobulin). Our computational analysis showed that the majority of interactions were between mouse monocyte-macrophage ligands and tumor epithelia, a finding potentially related to the high frequency of these cells in mouse tumors. While more complex procedures including spatial single cell RNA sequencing have recently established [29], immunopathological approaches that assess localization and distribution provide a cost effective and rapid means to quantitatively and qualitatively assess TIM-TIL localization in tumor sections. Such data may help differentiate poorly infiltrated tumors that contain high numbers of T cells from well infiltrated tumors with fewer immune cells.

We have identified certain limitations to our study. *First*, the diversity in the transcriptional landscapes observed between tumors could be related to the cell autonomous tumor. Understanding this relationship may result in the identification of additional TIM-TIL clusters or even immune cell types that associate with specific cell autonomous tumor lineage [11] or genomic landscapes. However, given the complexity of scoring clinical variables that impact the TIM status for each patient, a large number of clinical samples would be needed to draw causative relationships to any individual factor or feature of the CD45-negative tumors. Alternative approaches using lineage specific or mutationally defined mouse models may help delineate the effects of the cell autonomous tumor on CD45 immune cell infiltration. *Second*, our study included one type of bladder cancer mouse model. It will be important to determine whether species conservation exists between human and mouse tumors immune cells regardless of how tumor initiation occurs (e.g. genetic lesion versus carcinogen). Such analysis may provide important information related to the utility of different bladder cancer models for preclinical treatment studies.

Overall, we have identified gene expression programs found in infiltrating immune cells of primary human and mouse bladder tumors including the analysis of myeloid and lymphocytic cell types. Transcriptional analysis of single cell human and mouse bladder cancers highlighted both species conservation and transcriptomic divergence. These findings provide rationale to use the identified TIM-TIL populations and gene signatures for further progression and treatment related studies. Potential areas of interest include understanding patient TIM-TIL signatures that associate with *de novo* or acquired resistance to T cell targeted therapies and other TIM populations. Understanding that the BBN bladder model recapitulates certain key aspects of human tumor signatures, provides motivation to use this model to identify strategies that may increase the response rates to immune checkpoint inhibition.

## EXPERIMENTAL MODELS AND SUBJECT DETAILS

### Human sample acquisition

In this study we analyzed 10 human primary bladder cancer samples (Table S1). Samples were obtained from patients who underwent either trans urethral resection bladder tumor (TURBT) or radical cystectomy. Patient blood was drawn independently from the surgical procedure. This study was conducted under approval of the Mount Sinai School of Medicine IRB (10-1180, William OH) and written consent from all subjects. Human tissues samples were deidentified and subsequently not considered to be human subjects research. Resected tumor samples were examined by a genitourinary pathologist who confirmed the pathological staging and tumor content. Patient information is included in **Table S1**.

### Mouse tumor model

FVB/NJ mice (JAX, 001800) were treated with the 0.05 % OHBBN carcinogenic in drinking water for 14 weeks followed by 4 weeks of progression. Tumors at this stage consisted of muscle invasive disease and heterogeneous patterns of pathology and lineage composition. Age and sex matched tumors were used for transcriptomic analysis in this study. IACUC guidelines for proper animal husbandry and experimental use were followed (protocol LA13-00060).

### scRNA-seq of mouse and human bladder samples

Human and mouse tumors were then resected and dissociated to the single cell level in a solution of RPMI-10% FBS (Gibco) + 1% type 1 collagenase (Gibco). The dissociate is subject to mechanical dissociation through a 16G needle and syringe (5-10 passes) and then a 21.5G needle (5-10 passes). Single cell dissociates were then sorted for live CD45 positive cells and sorted using a BD Aria after cell staining using a BV510-CD45 antibody (Biolegend) and DAPI stain. Single cell suspensions of all sorted samples were then resuspended in BSA (0.2 % in PBS) at a concentration of 2×10^6^ cells/ml followed by barcoding with a 10x Chromium Controller (10x Genomics).

## SC-RNA SEQUENCING AND DATA PROCESSING

### Data pre-processing

R package *Seurat* [30, 31] was used to process both human and mouse scRNA-seq data. Cells with more than 2500(5000) genes detected were removed from human(mouse) scRNA-seq samples for concerns of doublet contamination. Different cutoffs were used for human and mouse data to keep the percentage of doublets (6.5%) similar between human and mouse. In addition, cells with percentage of mitochondrial genes higher than 5% (10%) were removed from human(mouse) data. Again, the different cutoffs were chosen to keep the percentage of possible dying cells removed (10%) to be similar between human and mouse. Gene counts were normalized and log-transformed using *Seurat*. Batch effect among samples are removed by *Seurat* MNN algorithm [30, 31].

### Broad immune cell clustering and annotation

The top 2000 most variable genes were scaled with the effects of total UMI count and percentage of mitochondrial genes regressed out using *Seurat*, from which principal components (PCs) were calculated. The top 30 PCs were selected for SNN clustering [32]. The number of PCs were decided by *Seurat* function *ScoreJackStraw* [30, 31]. To choose the optimal number of clusters, we altered the resolution parameter and manually checked the resulting clusters and their top marker genes. The optimal cluster number was chosen where the top marker genes showed most consistency with the canonical markers (see below). Each cell cluster was manually annotated by comparing the top marker genes for that cluster and canonical markers for different immune cells: CD3/CD4/CD8 (T cell marker), NKG7 (NK cell marker), LYZ (monocyte/macrophage marker), CD1C (DC marker), MS4A1 (B cell marker), MZB1 (plasma cell marker), MS4A2 (mast cell marker), IL3RA (pDC marker), CD24 (neutrophil marker). Non-immune cell clusters including erythrocytes (marked by HBB), platelets (marked by GP9), epithelial (marked by KRT19), endothelial (marked by PLVAP), and fibroblasts (marked by MMP2), were removed from further analysis. The cell clusters were visualized in UMAP [31].

### Refined subset clustering within Monocyte/Macrophage and T-NK cluster

The cells within the cluster of monocytes/macrophages from the previous immune cell type clustering procedure were extracted for further analysis. The top 2000 most variable genes within these monocytes/macrophages were used for calculating PCs and clustering following the same procedure as described above. The cell clusters were annotated as monocyte by highly expressed S100A4/S100A6/S100A8/S100A9 or macrophage by highly expressed C1QA/C1QB/C1QC. Different monocytes or macrophages subsets were annotated by top over-expressed genes for each cluster as calculated by *Seurat* function of *FindAllMarkers* [30, 31]. Similarly, cells within the T-NK cell cluster were further clustered into subsets using the above procedure. T cell subsets were annotated according to the classical cell markers, i.e., CCR7 (naïve), LAG3 (exhausted) and FOXP3 (regulatory), or top over-expressed genes in each subset.

### Immune cell type proportion

For each human and mouse sample, the proportion of each broad immune cell types were calculated. (Non-immune cells were excluded here.) To test the difference of the cellular proportion between human and mouse, both Wilcoxon rank-sum test and Student’s t-test were used.

### Comparison of expression profiles of cell subsets

To compare the gene expression profile of T-NK subsets between human and mouse, we followed similar procedures used in the previous study [9]. Briefly, a list of variable genes (defined below) across T-NK subsets were obtained for human and mouse datasets separately. Assuming *g_ij_* annotated the average expression (before log-transformation) of gene *i* in cell set *j*, we sorted *g_ij_* for each gene among cell sets of interests (i.e., T-NK cells):g_i1_<g_i2_<…<g_iN_, and those genes with max (g_iN_/g_iN-1_, …, g_i3_/g_i2_, g_i2_/g_i1_) > 1.2 were defined as variable genes across these cell sets. The *g_ij_* was then scaled gene-wisely across T-NK cells for human and mouse dataset separately. Common variable genes between human and mouse datasets were used to cluster and compare the expression profiles of T-NK cell sets. In heatmaps, hierarchical clustering was used with Pearson’s correlation coefficient as similarity measurement and complete linkage as the clustering method. The comparison of macrophage/monocyte expression profiles between human and mouse were conducted in the similar way, as well as that between bladder and lung cancer.

### Computational cell-cell interaction analysis

Interaction scores are defined as multiplication of average expression of ligand and average expression of receptor [19]. For calculation of pooled interaction scores, expressions are averaged from all samples. All interactions with one score bigger than 4 in at least one pair of cell types are of interest. Interaction scores of individual samples are also calculated. P-values of t-test by comparing interaction scores of individual samples and 0 are shown in heatmap. P-values of t-test by comparing scores of the same interaction and the same pair of cell types between human and mouse are shown in heatmap.

### Pathway activation analysis

Pathway activation is described by module score, calculated by R package *Seurat* function *AddModuleScore.* One module score is calculated for each of the pathways and each of the samples/clusters. To compare the activation of different samples, rank-sum test is performed on module scores.

### M1/M2 characteristics of clusters of Macrophages

M1 or M2 characteristics genes are set to be M1 module or M2 module, respectively. Module scores are then calculated by R package *Seurat* function *AddModuleScore* for each of the clusters. M1 and M2 module scores are compared by rank-sum test is performed on module scores to determine whether a cluster is M1 or M2 Macrophages.

### Immunohistochemical (Immunostat) analysis of immune cell localization

To assess immune cell numbers and localization within bladder tumors, consecutive tissue sections were stained for using conventional immunohistochemistry methods and DAB exposure. Stained slides were then digitally scanned and imported Adobe Photoshop for conversion to pseudo coloring and image overlays. Channel merged images were then imported to QuPath software where cells were counted with respect to 4 defined regions of equal spacing with respect to the lesion center. Cell numbers were determined for each region by subtracting values from the preceding regions. For example, Region 3 cell numbers = Region 3 ROI counts – Region 3 ROI counts.

## Acknowledgements

We acknowledge the Human Immune Monitoring Center at Mount Sinai.

## Author Contributions

HY: concept design, bioinformatic analysis and manuscript writing. JS: concept design, supported human scRNA sequencing and manuscript writing. LW: concept design, bioinformatic analysis and manuscript writing. JD: animal husbandry, immune cell isolation and manuscript review. MG: manuscript review and supported human scRNA sequencing. NB: manuscript review and supported human scRNA sequencing. OE: manuscript review. BF: manuscript review. JZ: provided support to HY and manuscript review. DM: wrote the manuscript and supported all aspects of the study.

## Competing Interests

No competing interests are reported.

## SUPPLEMENTAL INFORMATION

**Fig. S1. QC analysis for isolated CD45-negative and CD45-positive cellular fractions used for scRNA-seq**. (A) Human and (B) mouse QC dot plot analysis for RNA (left) and mitochondria (right) quality control.

**Fig. S2. scRNA-seq analysis of human and mouse CD45-negative tumor cells**. (A) Human and mouse merged UMAP clusters for CD45-negative cells. (B) Heat maps comparing human CD45 negative cells (left) and CD45 negative cells from mouse primary tumors (middle) and a mouse subcutaneous tumors (right). (C) UMAP clustering of mouse CD45-negative tumor cells by sample (left), epithelial source (middle), epithelial cluster and human epithelia by sample (far right). (D, left) UMAP clustering by sample for CD45-negative human (left) and mouse tumors (right). (D, inset and right). Combined human and mouse UMAP clustering for CD45-negative tumor cells and zoom inset (right). Human tumor 171 is more similar to mouse subcutaneous tumor 3928 while human tumor 2a is more similar to mouse tumors 8524 and 8525.

**Fig. S3. Distribution of NK and T cells in mouse and human bladder tumors**. (A) Left, distribution of T-NK cells in human (n=10) and, right, mouse bladder tumors (n=3). (B) Dot plots showing distributions of T-NK cell populations in human (black) and mouse (red) independent tumor samples. (C) Box plots showing distribution of major immune cell types in human and mouse tumors.

**Fig. S4. Comparison of human and mouse T-NK cells**. (A) Clustering for similarities between T-NK cells in human and mouse bladder tumors. (B) Heatmaps showing gene expression patterns in T-NK cell populations in human and mouse immune cell populations.

**Fig. S5. Clustering analysis for monocytes and macrophages in human and mouse bladder tumors**. UMAP clustering, heatmap analysis and representative genes for human monocytes and macrophages in human (A-C) and mouse (D-F) bladder tumors.

**Fig. S6. Clustering for species similarities in macrophage-monocyte gene expression between species and between organs**. Gene expression alignments between (A) human *versus* mouse bladder, (B, top) human lung *versus* human bladder, *bottom*, mouse lung *versus* mouse bladder and (C) similarities between macrophage-monocyte populations within immune cell populations for human (C, left) and mouse (C, right).

**Fig. S7. Pathway enrichment analysis in monocyte-macrophages populations in human and mouse bladder tumors**. (A) Comparison of various signaling pathways between monocyte-macrophage populations. (B) Identification of outlier cell populations in MYC, PI3K-AKT, interferon-gamma and TNF-alpha signaling between each pair of monocyte-macrophage populations. Module scores for each human or mouse subcluster were used to get p-values from rank-sum tests to determine whether there is a difference of pathway activation between pairs. See **Supplemental Data 5 and Data 6**.

**Fig. S8. Expression of genetic and immune targets in various immune cell populations in (A) mouse and (B) human bladder immune cell populations**. High expression of *XCR1-Xcr1* in human and mouse DCs only. High expression of *CD8A-Cd8a* in human T-NK cells and mouse pDCs. High expression of *CCR2-Ccr2* in human but not mouse monocytes-macrophages. High expression of *S100A8-S100a8* in human but not mouse monocytes-macrophages. High expression of *CD207-Cd207* in both human and mouse DCs. Low expression of *NR4A1-Nr4a1* in human and mouse monocytes-macrophages only.

**Fig. S9. Single cell expression for immune checkpoint proteins for individual human and mouse bladder tumors**. (A) Single cell expression of *PDCD1-pdcd1*, *HAVCR2-havcr2*, *CTLA4-Ctla4*, *CD274-Cd274* and *LAG3-Lag3* for human bladder patients (n=10) and mouse patients (n=3). (B) heat maps showing comparison between different immune blockade proteins. Those having greater red expression are more divergent according to Wilcox p values.

**Fig. S10. Application of a pathology based immunostat to measure immune cell numbers and localization in bladder tumors**. (A) Immunostains for T cells, NK cells and B cells shown as DAB exposure and after pseudo color conversion. (B) Example of how immune cells are quantitatively and qualitatively evaluated with respect to the lesion perimeter.

## TABLES

**Table 1**. Clinical and demographical information for human bladder tumors.

**Table 2**. Cell numbers mean reads and genes detected for individual human and mouse bladder tumors.

## SUPPLEMENTARY DATA FILES

**Supplementary Data 1**. Correlation between human and mouse epithelial clusters in bladder tumors

**Supplementary Data 2**. Pathway analysis for primary mouse tumors normalized to subcutaneous bladder tumors.

**Supplementary Data 3**. Correlation between human and mouse NK and T cell populations.

**Supplementary Data 4**. Correlation between human monocytes and macrophages.

**Supplementary Data 5**. Correlation between mouse monocytes and macrophages.

**Supplementary Data 6**. Hallmark pathway values for human monocytes and macrophages.

**Supplementary Data 7**. Hallmark pathway values for mouse monocytes and macrophages.

**Supplementary Data 8**. p-values for human *versus* human monocyte-macrophages populations.

**Supplementary Data 9**. p-values for mouse *versus* mouse monocyte-macrophages populations.

**Supplementary Data 10**. p-values for human *versus* mouse monocyte-macrophages populations.

**Supplementary Data 11**. p-values used to designate monocyte and macrophages populations as M1 or M2 in human bladder tumors.

**Supplementary Data 12**. p-values used to designate monocyte and macrophages populations a M1 or M2 in mouse bladder tumors.

